# An arylsulphonamide that targets cell wall biosynthesis in *Mycobacterium tuberculosis*

**DOI:** 10.1101/2024.07.22.604653

**Authors:** Renee Allen, Lauren Ames, Vanessa Pietrowski Baldin, Arielle Butts, Kenneth J. Henry, Diana Quach, Joseph Sugie, Joe Pogliano, Tanya Parish

## Abstract

We investigated the mechanism of action of an arylsulphonamide with whole cell activity against *Mycobacterium tuberculosis*. We newly synthesized the molecule and confirmed it had activity against both extracellular and intracellular bacilli. The molecule had some activity against HepG2 cells but maintained some selectivity. Bacterial cytological profiling suggested that mechanism of action was via disruption of cell wall synthesis, with similarities to an inhibitor of the mycolic acid exporter MmpL3. The compound induced expression from the IniB promoter and caused a boost in ATP production but did not induce reactive oxygen species. A mutation in MmpL3 (S591I) led to low-level resistance. Taken together these data confirm the molecule targets cell wall biosynthesis with MmpL3 as the most probable target.

---

Infections caused by *Mycobacterium tuberculosis* still remain a serious global health problem with >1 million deaths every year^1^. In order to improve drug treatment and overcome resistance, the pipeline of new drugs need to be constantly refilled. Phenotypic screening has identified many new drug classes with potential for development. Identification of the bacterial target for whole cell active compounds is an important component of hit evaluation and progression.

We are interested in identifying the targets of molecules identified in whole cell screens. An arylsulphonamide was identified by screening the AstraZeneca corporate library (320,000 compounds) and limited structure-activity relationship studies conducted^2^. The series appeared to have a novel target, since it did not inhibit mycolic acid synthesis (InhA), oxidative phosphorylation, DNA replication, transcription or translation, and the target was therefore uncharacterized^2^.

TPN-0157345 (2-methyl-4-(1*H*-pyrazol-1-yl)- *N*-(1-(pyridin-4-ylmethyl) piperidin-4-yl) benzenesulfonamide) was prepared according to the literature procedure.^2^ We found the inhibitory concentration (IC_90_) against aerobically cultured *M. tuberculosis* H37Rv was 3.3 μM which is consistent with the previously reported minimum inhibitory concentration (MIC) of 1.6 μM^2^ (Table 1). We also tested the molecule for activity against intracellular *M. tuberculosis* H37Rv in THP-1 macrophages and found the IC_90_ was 1.4 μM, again consistent with the previous report that the molecule had activity against bacteria inside THP-1 cells, although an MIC was not determined previously^2^. We determined cytotoxicity against the HepG2 liver cell line; the IC_50_ was 22 μM giving a selectivity index of ∼11. Previous work tested cytotoxicity against THP-1 cells, which was very similar at 32 μM^2^.

**Table 1.** Compound activity. Activity against *M. tuberculosis* strains (all derived from H37Rv London Pride (ATCC 25618) was measured in Middlebrook 7H9 medium with 10% v/v Middlebrook OADC supplement and 0.05 % w/v Tween 80 ^3^ or in THP-1 infected cells (MOI of 1)^4^. Cytotoxicity was measured against HepG2 cells^4^. Data are the average and standard deviation of two independent experiments. Fold change is between wild-type and MmpL3 mutant strains (IC_90_).

|  | IC<sub>50</sub> (µM) | IC<sub>90</sub> (µM) | Fold-change |
| --- | --- | --- | --- |
| Extracellular activity (wild-type) | 2.1 ± 0.2 | 3.3 ± 1.3 |  |
| Intracellular activity | 0.5 ± 0.06 | 1.4 ± 0.7 |  |
| HepG2 cytotoxicity | 22 ± 4.2 | ND |  |
| Extracellular activity<br>(MmpL3 F255L, V646M, F644I) | 1.4 ± 0.8 | 5.6 ± 5.3 | 1.7 |
| Extracellular activity<br>(MmpL3 S591I) | 4.1 ± 0.7 | 9.2 ± 2.5 | 2.8 |

Once we had confirmed that the newly synthesized molecule was active and retained similar properties to the previous report, we conducted bacterial cytological profiling (BCP) to identify the pathway(s) that are inhibited by this molecule^5^. BCP uses changes in cellular morphology effected by compound exposure to classify target pathways. *M. tuberculosis* mc^2^6206 was exposed to compounds for 48 and 120 h at two concentrations (Figure 1). The bacterial cells became rounded or ovoidal with increased membrane permeability at both concentrations. Comparison with a database of control molecules revealed the closest similarity to bacteria treated with a known MmpL3 inhibitor, the adamantyl urea AU1235, suggesting that the compounds affect the cell wall.

**Figure 1.**
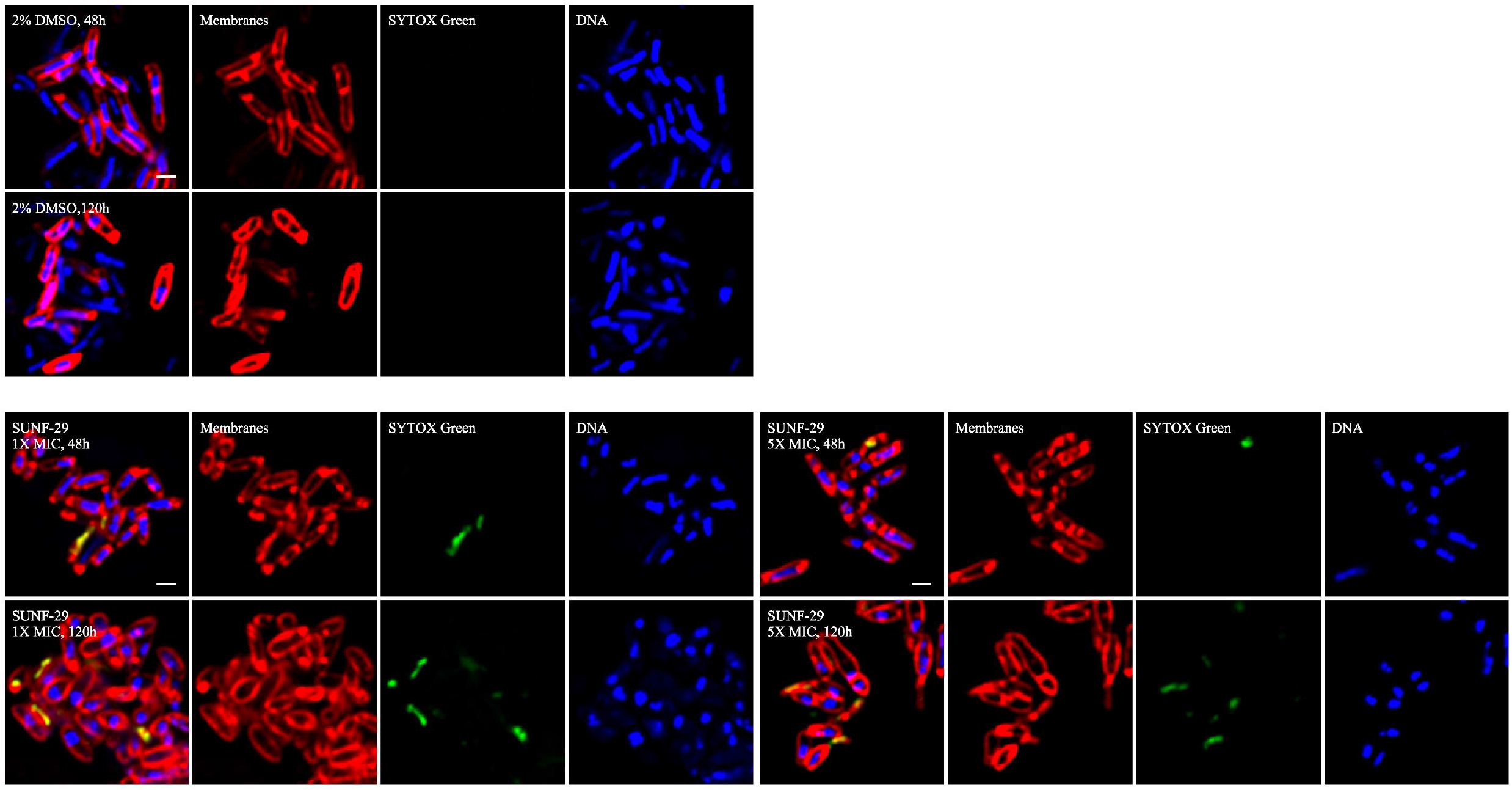
Bacterial cytological profiling. *M. tuberculosis* mc^2^6206 was inoculated into medium at an OD of 0.06 to 0.08, treated with compounds for 48-120 h and fixed with 16% paraformaldehyde, 8% glutaraldehyde and 0.4 M phosphate buffer (pH 7.5) Cells were washed twice with warm medium and stained for 30 min with FM 4-64 (Invitrogen), SYTO 40 (Invitrogen) and SYTOX Green (Invitrogen). Membranes are imaged in red, DNA in blue, and green staining indicates that cell integrity has been compromised; scale bars are 1 μm.

Our data suggested that the molecule targets cell wall synthesis. Increased expression of the IniBAC operon is observed with inhibitors of cell wall synthesis including MmpL3 inhibitors^6^. Therefore, we looked at induction of the IniB promoter using a reporter strain of *M. tuberculosis* (Figure 2). We saw a clear concentration-dependent increase in expression from P_iniB_ at concentrations around the MIC (at higher concentrations the induction is lost, presumably due to cell death). Similarly, the positive control, ethambutol, led to increased expression from P_iniB_ in a concentration-dependent fashion.

**Figure 2.**
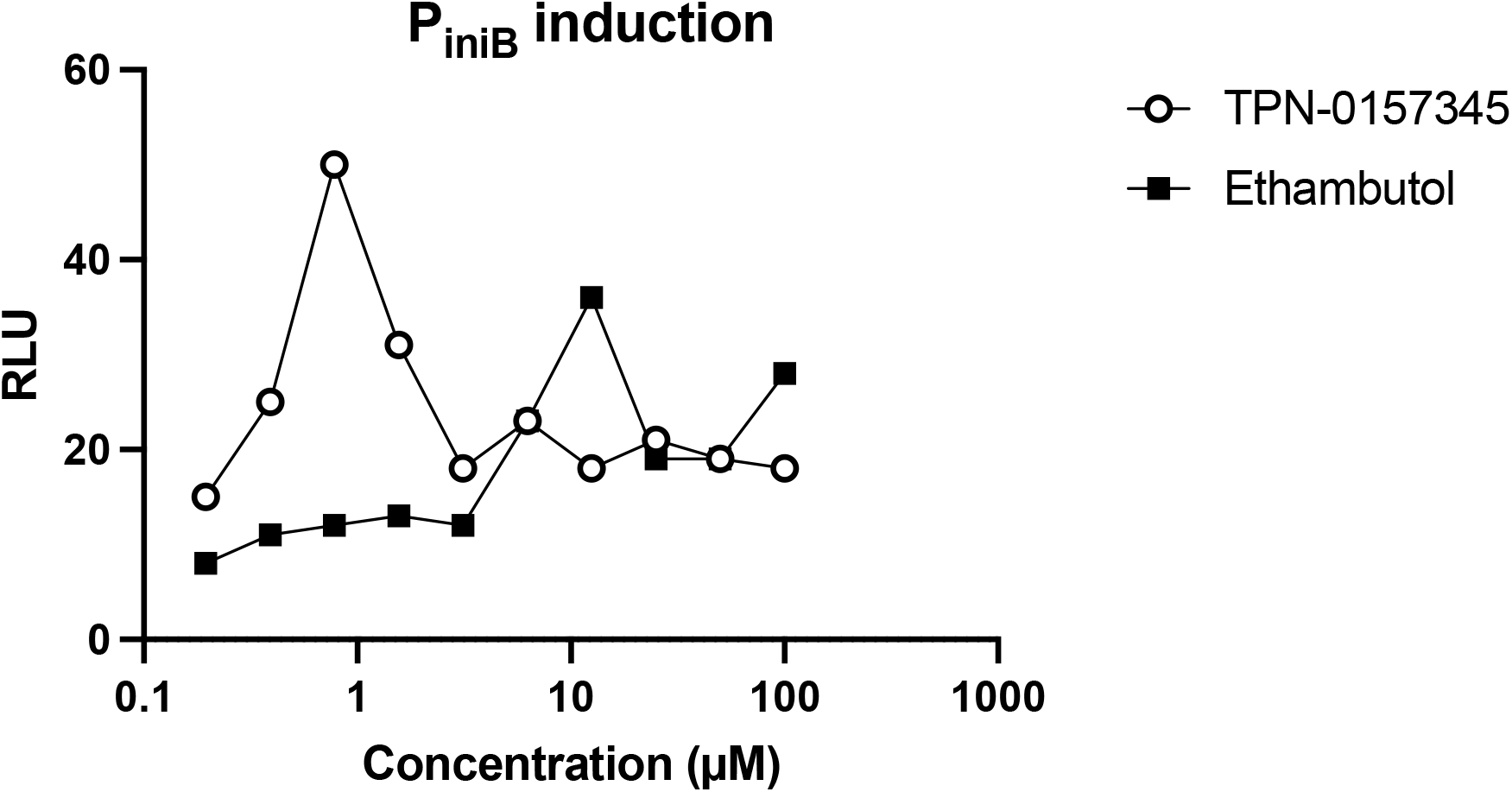
Exposure to the acylsulphonamide induces PiniB in *M. tuberculosis*. Recombinant *M. tuberculosis* expressing Lux under the control of the IniBAC promoter (P_iniB_-Lux)^6^ was cultured to logarithmic phase and exposed to compounds for 3 days. Luciferase activity was measured after the addition of luciferin by measuring relative luminescence units (RLU). Data are representative of two fully independent runs (See Figure S1).

We tested whether compounds induced a general stress by determining whether there was any generation of reactive oxygen species (ROS) (Figure 3). We used the fluorescent reporter dye dichlorodihydro-fluorescein diacetate to monitor of ROS. No generation of ROS was seen after exposure to the arylsulphonamide compound confirming that this is not the mechanism of action. The positive control, econazole, induced ROS in a concentration dependent fashion.

**Figure 3.**
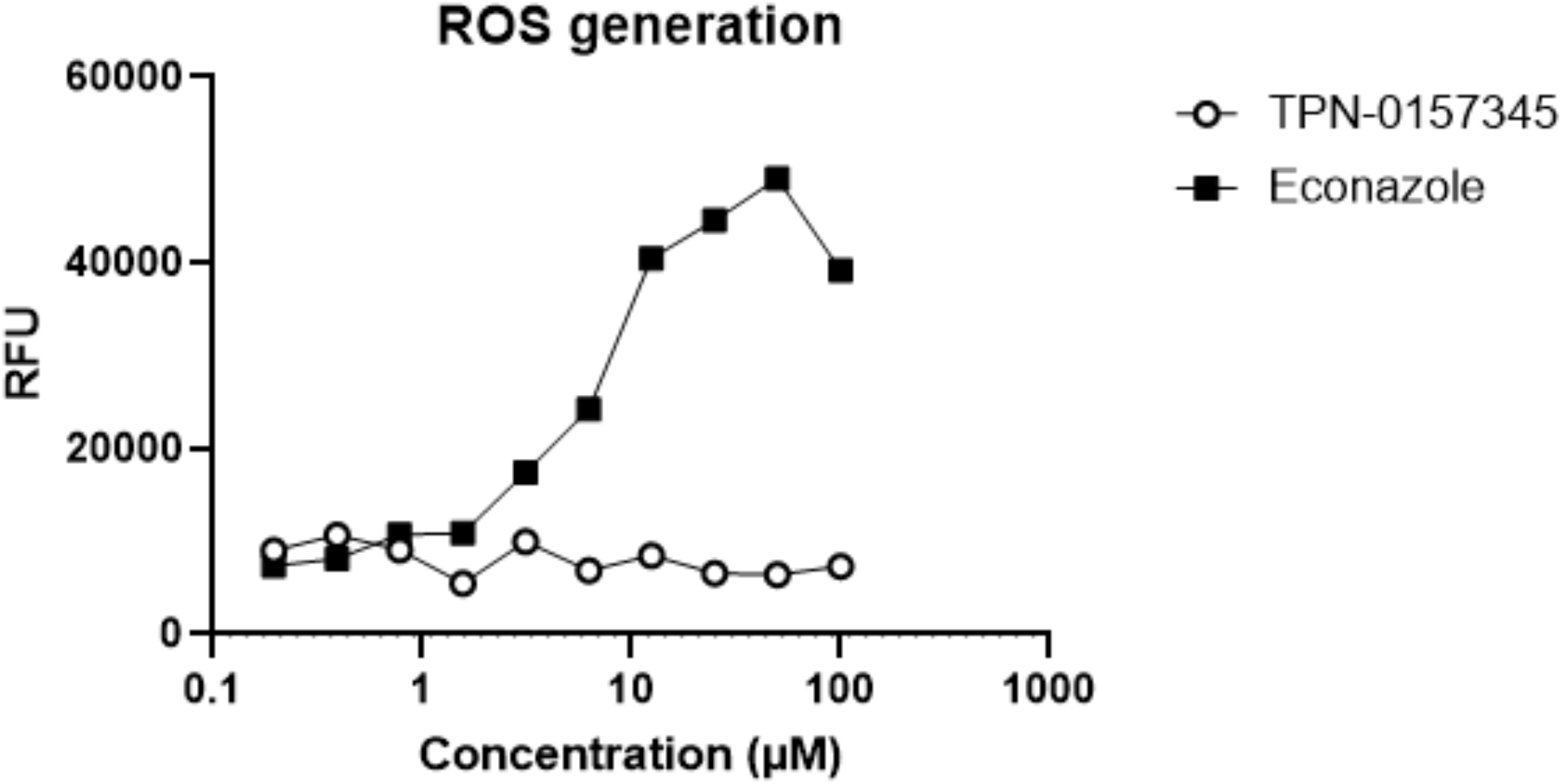
Exposure to the acylsulphonamide does not induce the production of reactive oxygen species in *M. tuberculosis*. *M. tuberculosis* (H37Rv) was cultured to logarithmic phase and loaded with 40 μM dichlorodihydrofluorescein diacetate. Cultures were exposed to compounds for 90 min and the production of ROS was determined by measuring relative fluorescence units (RFU) at Ex485/Em535. Data are representative of two fully independent runs (See Figure S1).

Previous work has shown that cell wall inhibitors can lead to a boost in ATP production in *Mycobacterium bovis* BCG^7^. We tested the effect of the compound on ATP levels. We saw an increase in ATP levels at concentrations around the MIC (Figure 4). ATP levels remained high even when bacterial growth was inhibited, demonstrating increased ATP production consistent with targeting cell wall biosynthesis. As expected, the positive control Q203, which inhibits the electron transport chain, resulted in ATP depletion at concentrations below the MIC.

**Figure 4.**
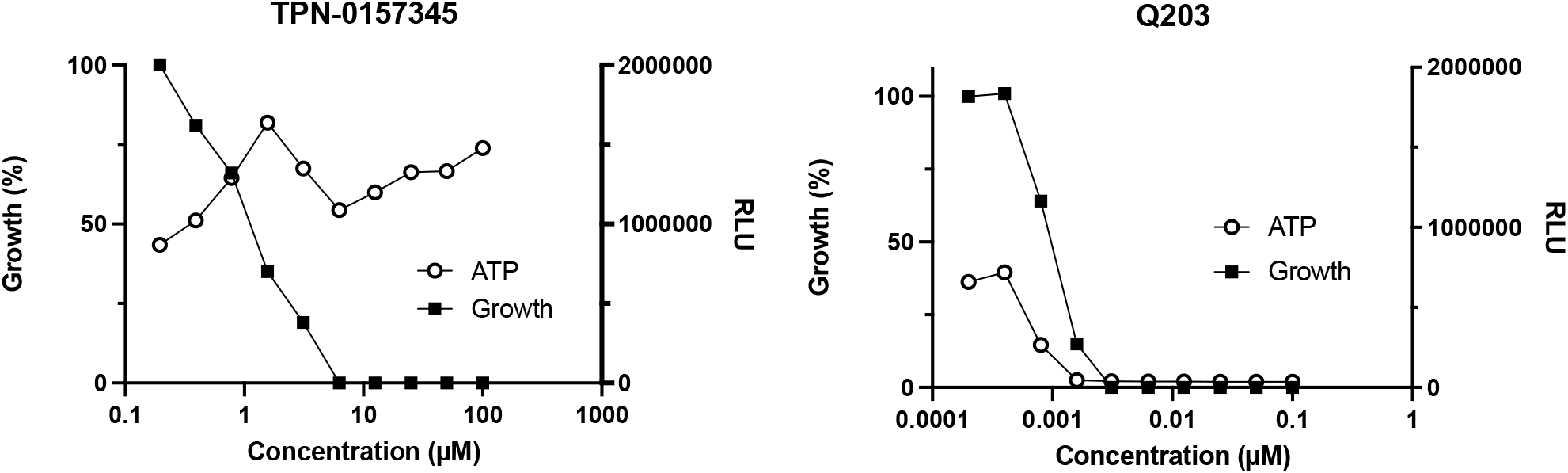
Exposure to the acylsuplhonamide leads to a boost in ATP production in *M. tuberculosis*. *M. tuberculosis* (H37Rv) was cultured to logarithmic phase and exposed to compounds for 24 h. ATP was measured using the BacTiter Glo reagent after incubation for 10 min in the dark and measuring RLU. Growth was determined after 5 d by reading OD_590_ and normalized to the DMSO control. Data are representative of two fully independent runs (See Figure S1).

MmpL3 is a promiscuous target and can be inhibited by many chemical scaffolds^8^. We tested whether the compounds were active against *M. tuberculosis* strains carrying mutations in MmpL3. We first tested a strain with three MmpL3 mutations (F255L, V646M, F644I)^9^ which is resistant to multiple chemical scaffolds but found no difference in MIC (Table 2). However, when testing a strain with a different (single) mutation (S591I), we saw a shift towards low-level resistance. These data are consistent with the molecule targeting MmpL3 but indicate that its mode of binding may be different from several chemical classes such as the spiral amines and adamantly ureas^9^.

In summary, exposure to the arylsulphonamide molecule inhibits cell wall synthesis and induces an ATP boost. Mutation in MmpL3 confers resistance to the molecule. Our data are consistent with a mode of action in which the arylsulphonamide targets MmpL3.

## Acknowledgements

We thank Grace Liu, Do Teon Park, Quan Pham and Felipe Santana for technical assistance.

